# H2-O deficiency promotes regulatory T cell differentiation and CD4 T cell hyperactivity

**DOI:** 10.1101/2023.08.14.553240

**Authors:** Robin A. Welsh, Nianbin Song, Chan-su Park, J. David Peske, Scheherazade Sadegh-Nasseri

## Abstract

Regulatory T cells (Treg) are crucial immune modulators, yet the exact mechanism of thymic Treg development remains controversial. Here, we present the first direct evidence for H2-O, an MHC class II peptide editing molecular chaperon, on selection of thymic Tregs. We provide evidence that lack of H2-O in the thymic medulla promotes thymic Treg development and leads to an increased peripheral Treg frequency. Single-cell RNA-sequencing (scRNA-seq) analysis of splenic CD4 T cells revealed not only of an enrichment of effector-like Tregs but also of activated CD4 T cells in the absence of H2-O. Our data support two concepts; a) lack of H2-O expression in the thymic medulla creates an environment permissive to Treg development and, b) that loss of H2-O drives increased basal auto-stimulation of CD4 T cells. These findings can help in better understanding of predispositions to autoimmunity and design of therapeutics for treatment of autoimmune diseases.

## Introduction

T cells are key players in humoral immune responses. Upon infection with a pathogen, CD4 T cells utilize their T cell receptor (TCR) to survey peptides presented by professional antigen presenting cells (APCs) on class II molecules (pMHCII) for recognition. Identification of a cognate pMHCII complex by the TCR leads to CD4 T cell activation and ultimately clearance of the foreign pathogen. Faulty activation, however, can lead to deleterious inflammation causing possible autoimmune diseases and cancer development. Therefore, multiple regulatory processes exist to ensure proper T cell activation.

Regulation of T cell activation begins during thymic development where immature thymocytes are screened for self-reactivity. Broadly divided into positive and negative selection, this process ensures that CD4 T cells expressing high avidity self-reactive TCRs are either eliminated (1), or become CD4 regulatory T cells (Tregs) (2). Critical to the screening process for identifying auto-reactive T cells is the Class II antigen processing machinery expressed by medullary thymic epithelium cells (mTECs) and thymic antigen presenting cells (APCs). Two chaperone proteins, H2-M (murine; human, HLA-DM), and H2-O (murine; human, HLA-DO) are major components of the MHC II processing pathway. While H2-M is expressed in all APCs, H2-O was originally discovered to be expressed in the thymic medulla and B cells (3). It was only later that HLA-DO was reported to be expressed on cross presenting CD8αα+ DC (4), then followed by identification of H2-O on murine CD8αα+ DC (5). H2-M plays a critical role in MHC class II antigen processing by dissociating the Class II Invariant Chain peptide (CLIP) from the newly synthesized MHC II. Dissociation of CLIP promotes a peptide-receptive MHC II conformation that can readily scan denatured antigens for the best MHC II groove fitting epitopes. However, a peptide-receptive MHC II conformation is highly transient (6, 7, 8, 9). We have proposed that H2-O binds to H2-M (10) and works cooperatively with H2-M by stabilizing the peptide-receptive MHC II conformation in order to optimize epitope selection (11). Together, these two molecules can ensure that the best MHC II groove fitting epitopes are presented to cognate CD4 T cells.

While the exact mechanism of Treg selection remains to be fully understood (12), two critical requirements have been identified as necessary for successful thymic Treg development. First, thymic Treg development requires relatively strong TCR signaling in the thymic medulla (13), and second, Treg development relies on signaling by common γ chain (γC) cytokines, mainly IL-2, for driving Foxp3 expression (14). Considering that strong TCR signaling during negative selection normally leads to CD4 T cell deletion (1), when investigating Treg selection, both the nature and density of medulla presented self-peptides must be considered. If epitopes are in high abundance and more ubiquitously expressed, then cognate CD4 T cells will undergo clonal deletion. However, if epitopes are in lower abundance and have a sparser expression, leading to discontinuous TCR stimulation, then cognate CD4 T cells might undergo Treg selection (12, 15, 16). This model of Treg selection relies on the level of TCR signaling that medulla localized CD4 single-positive (SP) T cells receive. Should a lower density of self pMHCII be presented in the thymic medulla to CD4 T cells undergoing negative selection, an increased number of self-reactive CD4 T cells might escape deletion, leading to an increased frequency of auto-reactive T cell clones in the periphery. Alternatively, presentation of a lower density of self pMHCII could promote selection of regulatory T cells. Here, we demonstrate that loss of H2-O generates a more stimulatory *in vivo* environment impacting both the thymic development and peripheral activation of regulatory T cells.

## Results

### Loss of H2-O increases the activation state of auditing medulla CD4 T cells

Previously, we demonstrated that loss of H2-O expression correlated with both an increased B cell presentation of low-affinity MHC II peptides, and an increased frequency of a MOG_35-55_ specific, self-reactive CD4 T cell (17). Because of H2-O expression in the medullary thymus we speculated that H2-O deficiency might lead to presentation of lower densities of high-affinity self-peptides in the medulla, thereby causing altered clonal deletion. Positively selected (TCR-β+ CD5+) medulla CD4 T cells (**Fig 1a/sFig1**) from the thymi of 6 week old male and female H2-O WT and H2-O mice were subdivided into “Auditing” (CCR7+ CD4+ Caspase-3_neg_) and “Clonally Deleted” (CCR7+ CD4+ Caspase-3_pos_) CD4 T cells for evaluation of negative selection.

**Figure 1:**
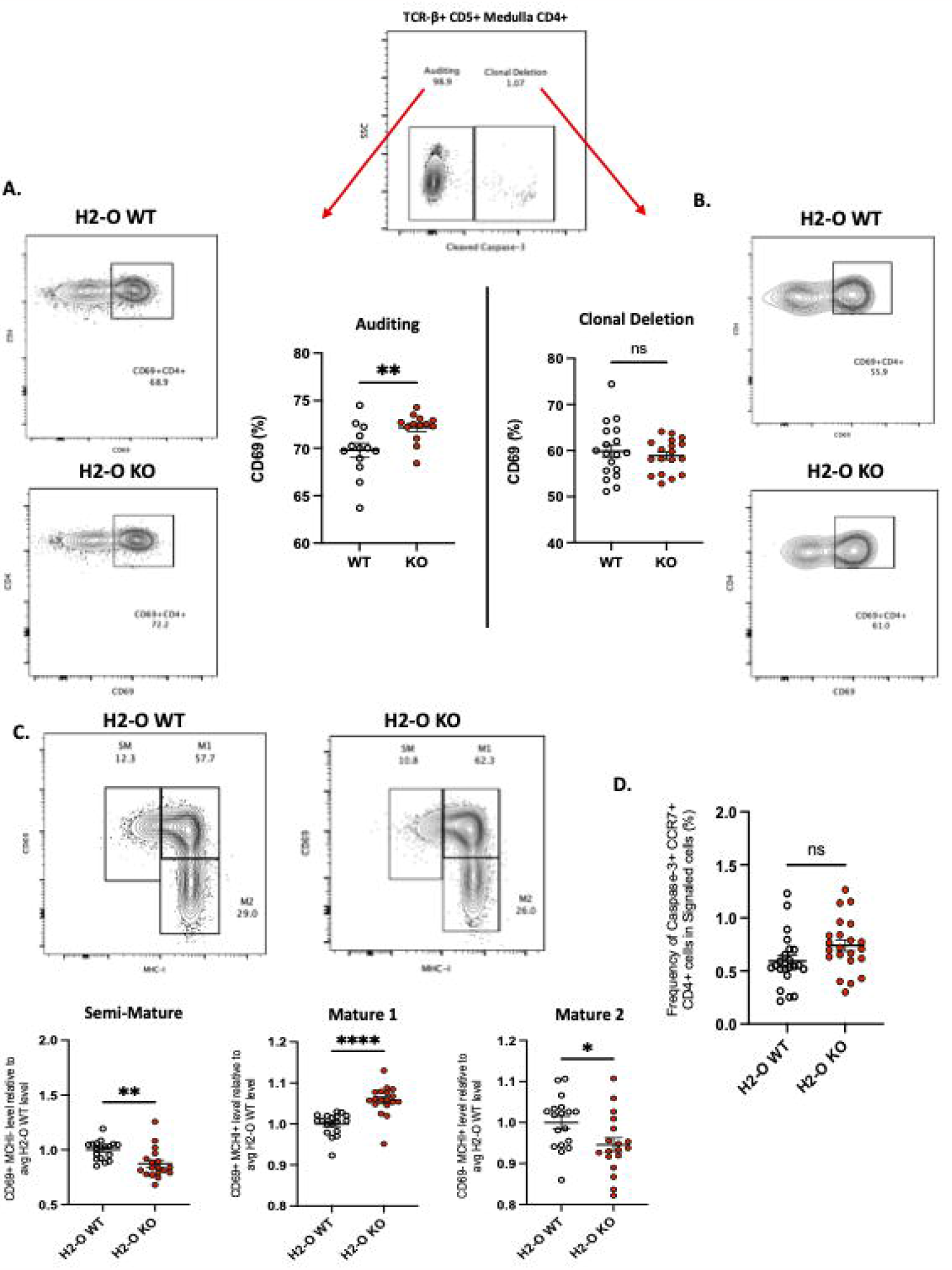
Loss of thymic H2-O increases the activation state of auditing CD4 T cells. **A. Left:** representative contour plots showing total CD69 expression in auditing (Caspase-3 negative) signaled (TCR-B+CD5+) CCR7+CD4+ T cells from 6-week-old H2-O WT (Top) and H2-O KO (Bottom) thymi. **Right:** Combined CD69 expression data from 5 repeat experiments. N= 18 mice per genotype **B. Right:** representative contour plots showing total CD69 expression in Clonally deleted (Caspase-3 positive) signaled (TCR-B+CD5+) CCR7+CD4+ T cells from 6-week-old H2-O WT (Top) and H2-O KO (Bottom) thymi. **Left:** Combined CD69 expression data from 5 repeat experiments. N= 18 mice per genotype **C. Top:** representative contour plots showing the subdivision of auditing (Caspase-3 negative) signaled (TCR-B+CD5+) CCR7+CD4+ T cells from 6-week-old H2-O WT (Left) and H2-O KO (Right) thymi into three maturation stages: Semi-Mature (SM), Mature 1 (M1), and Mature 2 (M2). **Bottom:** Cumulative maturation state data from 5 repeat experiments, N = 18 mice per genotype. Expression has been normalized to the average H2-O WT levels within each experiment to allow for comparison across experiments. Raw percentage data can be found in Supplemental Figure 3. **D**. Frequency of medulla specific (CCR7+) CD4 T cells selected for clonal deleted (Caspase-3+) ns = not significant * = <0.05 ** = <0.001 ***= <0.0001 **** = <0.00001 Statistics: unpaired student T-test

Loss of H2-O was found to significantly increase the expression of the activation marker CD69 on CD4 T cells undergoing active self-auditing, but not those selected for clonal deletion (**Fig 1a**). CD69 in combination with MHC-I defines 3 medullary maturation stages: semi-mature (CD69+ MHC-I -), mature 1 (CD69+ MHC-I +), and mature 2 (CD69-MHC-I -) (18). Subdivision of auditing H2-O-KO CD4 T cells identified a significant increase in CD4 T cells with a Mature 1 phenotype (**Fig 1b, sFig2**). Conversely, both the semi-mature and mature 2 stages were decreased in H2-O KO mice (**Fig 1b left/right**). No differences in any maturation stage were found in the clonally deleted CD4 T cell populations (**sFig2). Furthermore**, H2-O deficiency, did not appear to alter the rate of CD4 T cells undergoing clonal deletion (**Fig 1c**). Importantly, no differences were observed in thymocytes undergoing positive selection (**sFig 3**). These data suggest that loss of H2-O drives a more stimulatory thymic medulla environment, but with similar levels of clonal deletion. It is therefore likely that the increased peripheral frequencies of MOG-specific CD4 T cells previously identified is due to increased peripheral expansion of the MOG-reactive clone, not a general alteration in clonal deletion.

### H2-O KO thymi have increased frequencies of regulatory T cells

With increased levels of peripheral Tregs previously identified in H2-O KO mice (17), we also questioned if H2-O deficiency was affecting Treg selection. In fact, one model of thymic Treg selection centers around the concept of antigen density (15, 19). Within this “mosaic” model, sporadic MHC-TCR interactions with sparsely presented self-epitopes leads to Treg development. Since it has been shown that peripheral loss of H2-O leads to alterations in peptide presentation (17, 20, 21), we postulated that altering the level of self-epitopes present in the medulla could alter Treg selection.

Analysis of CD4 single-positive T cells identified an increased frequency of CD25+Foxp3+ T cells in H2-O KO mice (**Fig 2a**). Furthermore, H2-O KO thymic Tregs (tTregs) expressed higher levels of CD25, the high affinity IL-2 receptor, and Nur77, the orphan nuclear receptor (**Fig 2b**). As Nur77 has been associated with the level of TCR engagement (22), increased expression of Nur77 in H2-O KO Tregs suggests that they have been strongly signaled. Finally, maturation state analyses uncovered that H2-O KO tTregs were enriched in the M1 stage (**Fig 2c**).

**Figure 2:**
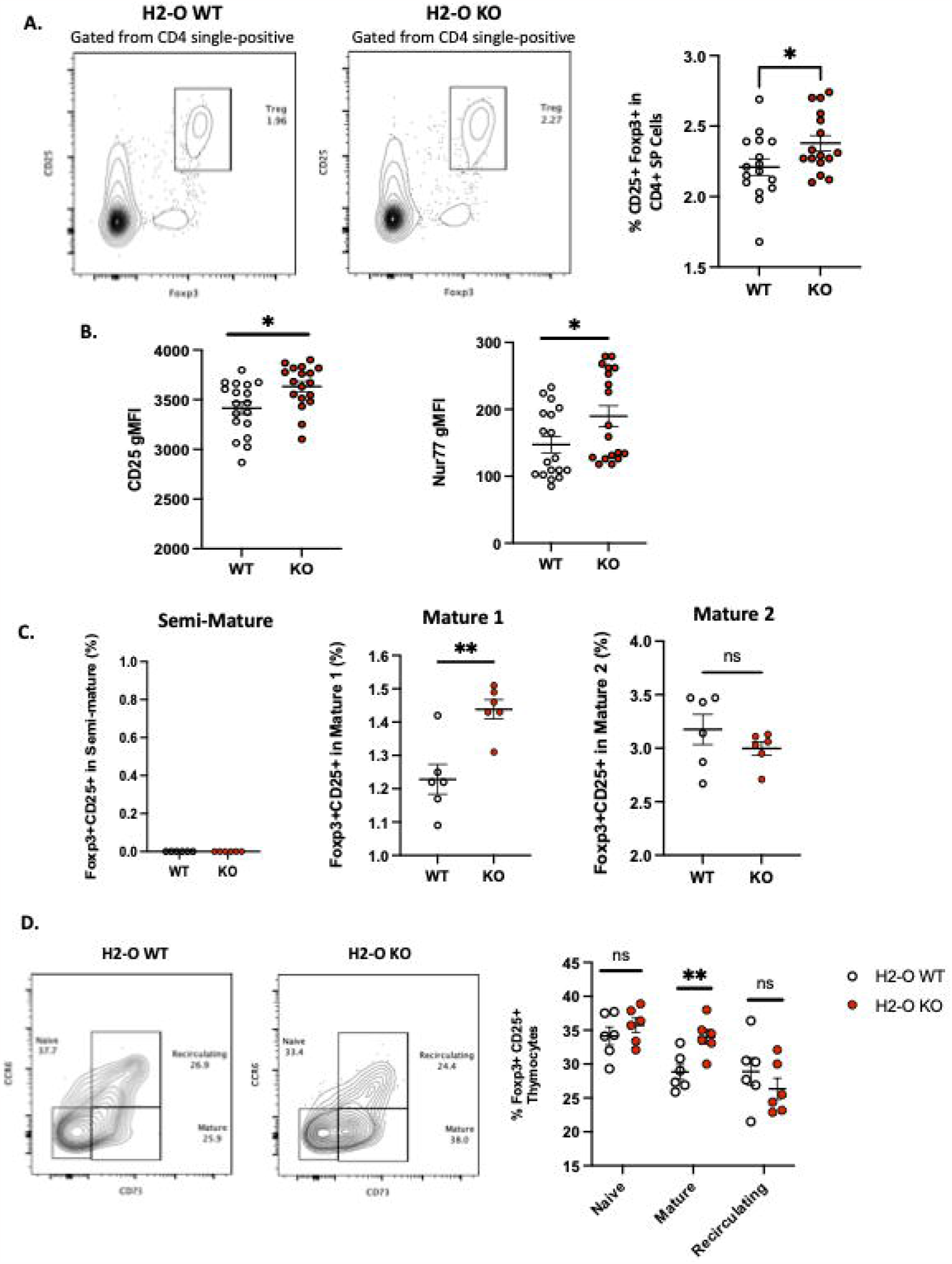
H2-O KO thymi have increased selection of regulatory T cells. **A.Left:** representative contour plots showing the frequency of Tregs (CD25+ Foxp3+) cells in the CD4 single-positive thymus population. **Right:** cumulative percentage of Foxp3+ CD25+ cells within the CD4 single-positive thymus population in H2-O WT (white) or H2-O KO (red) cells. Data from 3 replicate experiments, N= 16 mice per genotype. **B**.Geometric mean fluorescence intensity (gMFI) of CD25 (left), and Nur77 (right) expressed by Foxp3+ CD4+ T cells in the thymus of H2-O WT (white) or H2-O KO (red) mice. **C**.Subdivision of Treg cells into three maturation stages: Semi-Mature (Left), Mature 1 (Middle), and Mature 2 (Right). Data from 2 replicate experiments. **D.Left:** representative contour plots showing the frequency of Naive (CCR6-CD73-), Mature (CCR6-CD73+), and Recirculating (CCR6+ CD73+) Foxp3+ CD25+ Tregs in 6-week-old H2-O WT and H2-O KO thymi. **Right:** Summary plots showing the frequency of Naive (CCR6-CD73-), Mature (CCR6-CD73+), and Recirculating (CCR6+ CD73+) Foxp3+ CD25+ Tregs from 2 independent repeat experiments, N= 6 mice per group.

**Figure 3:**
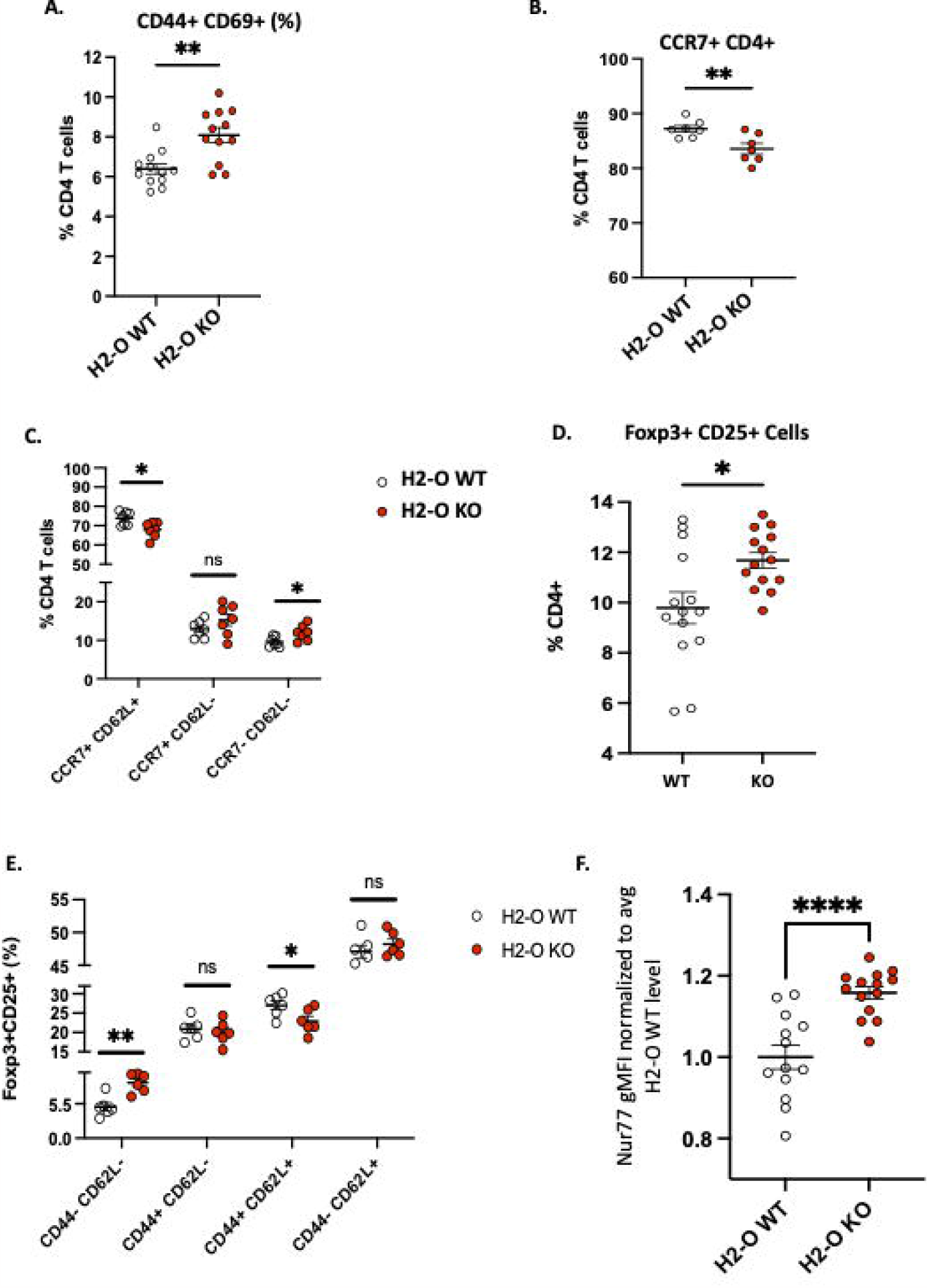
H2-O KO mice have an increased activated peripheral CD4 T cell population. **A**.Basal levels of CD44 and CD69 expressed by H2-O WT (white) and H2-O KO (red) splenic CD4 T cells. Increased CD69 expression, a marker of recent activation showed increased levels on H2-O KO CD4 T cells **B**.Percentage of CD4+ T cells expressing the lymphoid tissue homing receptor CCR7 **C**. CCR7 and CD62L expression levels in unimmunized splenic CD4 T cells **D**. Frequency of Foxp3+CD25+ cells in splenic CD4+ T cell population of unimmunized mice **E**. H2-O KO peripheral Tregs express decreased levels of naïve (CD44+ CD62L+) expressing cells **F**. Normalized Nur77 gMFI levels in Foxp3+CD25+ Treg cells in H2-O WT (white) and H2-O KO (red) cells. To account for experimental variation the average Nur77 gMFI level in H2-O WT samples was calculated. gMFI levels in both H2-O WT and H2-O KO samples were then divided by the calculated H2-O WT average. An increased Nur77 ratio indicates increased Nur77 gMFI levels. Summary of 3 repeat experiments. * = <0.05 ** = <0.001 *** = <0.0001 **** = <0.00001

Peripheral Tregs are known to recirculate back to the thymus (23, 24). We, therefore assessed the percentages of recirculating Tregs in H2-O Wt and H2-O KO thymi. **Figure 2d** shows that similar levels of recirculating (CCR6+ CD73+) Tregs were present in both H2-O WT and H2-O KO thymi. The results shown in Fig 2D are in agreement with the notion that increased Mature Tregs are of de novo origin rather than recirculating.

### Loss of H2-O correlates with increased peripheral CD4 T cell activation

Considering the observation that lack of H2-O did not appear to alter clonal deletion frequencies but did affect the level of thymic CD4 T cell activation, we next evaluated whether peripheral loss of H2-O also increased CD4 T cells activation. Unimmunized H2-O KO spleens had an increased frequency of CD4 T cells co-expressing the key activation markers CD44+ and CD69+ (**Fig 3a**) and the tissue homing marker CCR7 (**Fig 3b**). We further assessed the levels of “non-activated” (CCR7+ CD62L+) versus “activated” (CCR7-CD62L-) CD4 T cells (25), and found lower frequencies of non-activated CD4 T cells in H2-O KO mice (**Fig 3c, left**). Importantly, this correlated with an increase in percentage of activated CD4 T cells (**Fig 3c, right**). Collectively, these phenotypic analyses support the idea that loss of H2-O leads to increased basal levels of activated CD4 T cells in unimmunized H2-O KO mice.

As discussed above, H2-O KO thymi promoted Tregs selection. Consistent with these observations and our previously published data (17, 26), H2-O KO spleens had an increased frequency of CD25+ Foxp3+ Tregs (**Fig 3d**). Furthermore, H2-O KO Tregs showed decreased levels of CD62L (**Fig 3e**) and increased levels of Nur77 (**Fig 3f**) indicating these cells are likely more activated and circulating through the periphery.

### Single cell RNA-Sequencing of H2-O KO splenic CD4 T cells confirms increased activation

Based upon the strong FACS data above exhibiting increased numbers of Tregs, and more activated CD4 T cells, we attempted single-cell RNA-sequencing (scRNA-seq) to gain a more holistic unbiased characterization. CD3+ CD4+ NK1.1-CD19-cells were sorted from spleens of 3 unimmunized H2-O WT and 3 H2-O KO mice and subjected to 10x Genomics scRNA-seq analyses. In total, 11 distinct CD4 T cell clusters were identified (**Fig 4a & sFig4**). Separation of the clusters based upon H2-O WT or H2-O KO genotypes identified a dramatic increase in cluster 0, cluster 2 and cluster 3 in the H2-O KO samples. Conversely, clusters 1 and 4 were significantly increased in H2-O WT samples (**Fig 4b**).

**Figure 4:**
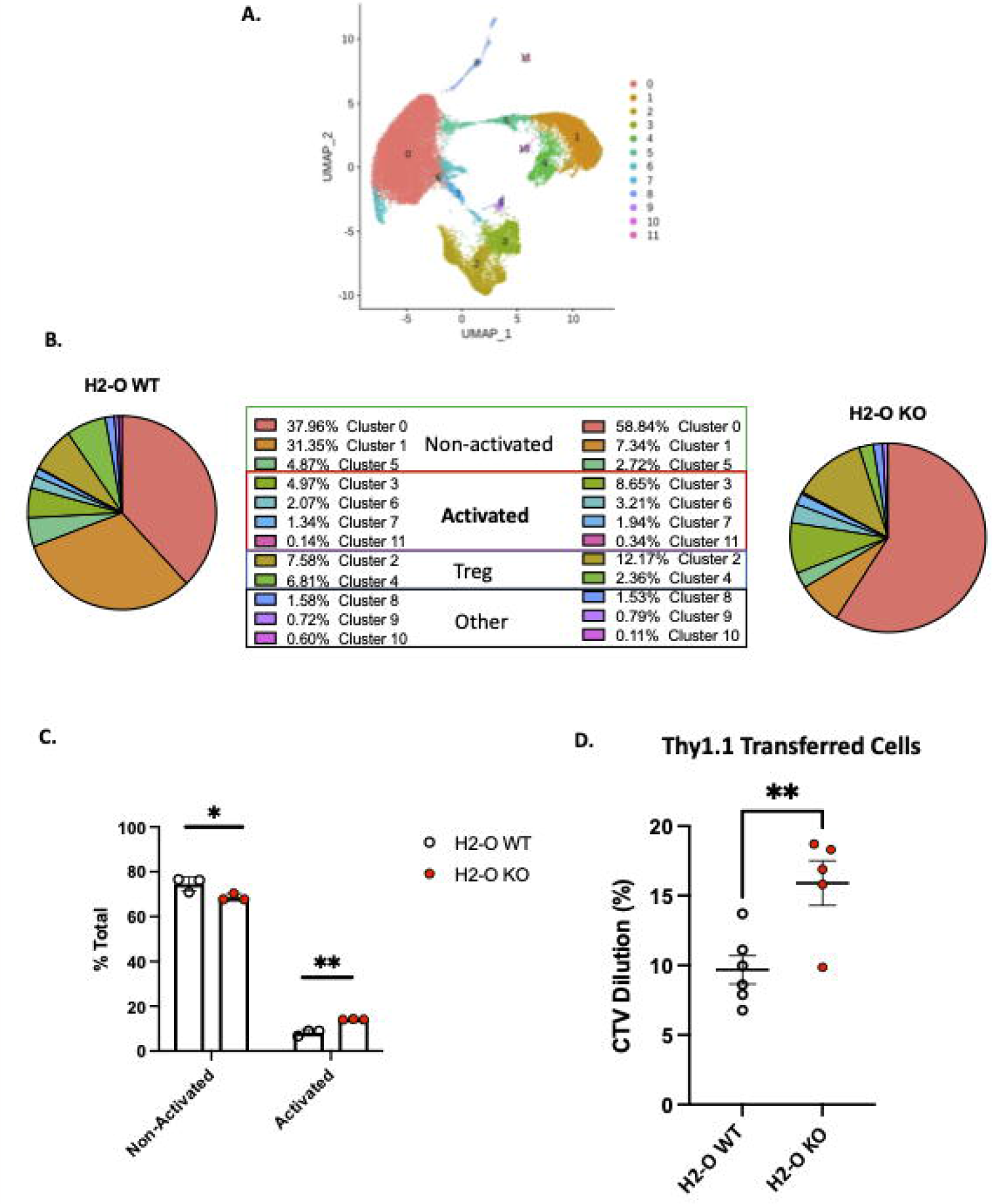
Loss of H2-O function causes increased basal CD4 cell activation. **A**.scRNA-seq clustering of CD4 T cells after Seurat analysis. Data represents the average of of 3 biological replicates per genotype. **B**.Breakdown of clusters in H2-O WT (Left) or H2-O KO (Right) samples. Clusters are grouped based upon, (1) known CD4 T cell subset markers and (2) gene comparison to published CD4 T cell data sets available on the Immunological Genome Project (www.immgen.org). Identified CD4 Cell phenotypes were: Non-activated, Activated, and Regulatory T cells. “Other” refers to a minor macrophage and NKT cell contamination from the sorting process. **C**.Distribution of the for the Non-activated and Activated phenotypes across the H2-O WT and H2-O KO biological replicates (N = 3 mice per genotype). **D**.In vivo proliferation of adoptively transferred naïve Thy1.1+ CD4 T cells (Thy1.1+ CD3+ CD4+ CD44-CD25-) after 7 days in either Thy1.2 H2-O WT (white) or Thy1.2 H2-O KO (red) hosts. Pooled data from 2 independent experiments (N= 6 H2-O WT, 5 H2-O KO)

Comparing the genes from Clusters 0, 1 and 5 to the published CD4 T cell datasets using the “My Geneset” function on the Immunological Genome Project (www.imgen.org) suggested that upregulated genes (Log_2_Avg Fold-change (FC) ≥0.5) were present in both “naïve” and “activated” CD4 T cell datasets. Based upon the high expression of CCR7 however, these clusters were labeled as “non-activated”. Clusters 3, 6, 7, and 11 aligned with mainly an “activated” CD4 T phenotype. Cluster 3 was found to have increased expression of the activation marker CD44, while Cluster 11 expressed high Ki67. Clusters 2 and 4 contained the known Treg genes Foxp3 and Il2ra (CD25) (**Fig 4a & Sup Table 1**). Clusters 8, 9, and 10 were small populations of Macrophages and NKT cell (**Sup Table 1**). Condensing the cluster analysis further supported our initial FACS data that unimmunized H2-O-KO mice have a decreased frequency of “Non-activated” CD4 T cells (68.90% avg H2-O KO vs 74.17% avg H2-O WT with a coinciding increase in “Activated” (14.14% avg H2-O KO vs 8.53% avg H2-O WT) CD4 T cells (**Fig 4b**).

To test whether the higher activated state in H2-O KO T cells were due to the H2-O KO CD4 T cells themselves, or possibly induced by the *in vivo* environment, we performed an adoptive transfer experiment of dye labeled naïve (CD4+ CD44-CD25-) Thy1.1+ CD4 T cells from H2-O WT into either unimmunized Thy1.2+ H2-O WT, or Thy1.2+ H2-O KO hosts. As shown in **Fig 4d**, H2-O KO hosts induced significantly more *in vivo* proliferation of the Thy1.1+ WT cells than the H2-O WT hosts. These data additionally support the idea that loss of H2-O creates a more activating CD4 T cell environment by presenting a wider range of self-epitopes (17).

### H2-O KO Tregs have an activated phenotype

Analyses of the two Treg clusters revealed that the frequency of Clusters 2 and 4 were roughly equal in H2-O WT samples, whereas H2-O KO samples had an overrepresentation of Cluster 2 (**Fig 5a, left**). Both cluster 2 and 4 expressed the classic Treg genes Foxp3, IL2ra [CD25], CTLA4, and Pdcd1 [PD-1] (**Fig 5a, right**; **Sup Table 1**). Also identified was the transcription factor Ikzf2 [Helios], a marker of stable Treg lineage commitment in inflammatory conditions (27) and debated marker of thymic Tregs (28, 29, 30, 31). Supporting our scRNA-Seq analysis, Helios protein expression was found in both H2-O WT and H2-O KO CD4 Tregs (**Fig 5b**). Comparison of the cluster 2 genetic signature with a published Treg data set indicated that these cells are likely effector-like Tregs. Furthermore, breaking down Cluster 2 by genotype showed that H2-O KO Tregs had a larger fold change in expression of the core effector genes signature (32) (**Fig 5c**). We hypothesize that this activated Treg state is likely driven by the increased basal CD4 T cell activation. Expansion of this Treg population could certainly be a mechanism controlling spontaneous autoimmunity in H2-O KO mice.

**Figure 5:**
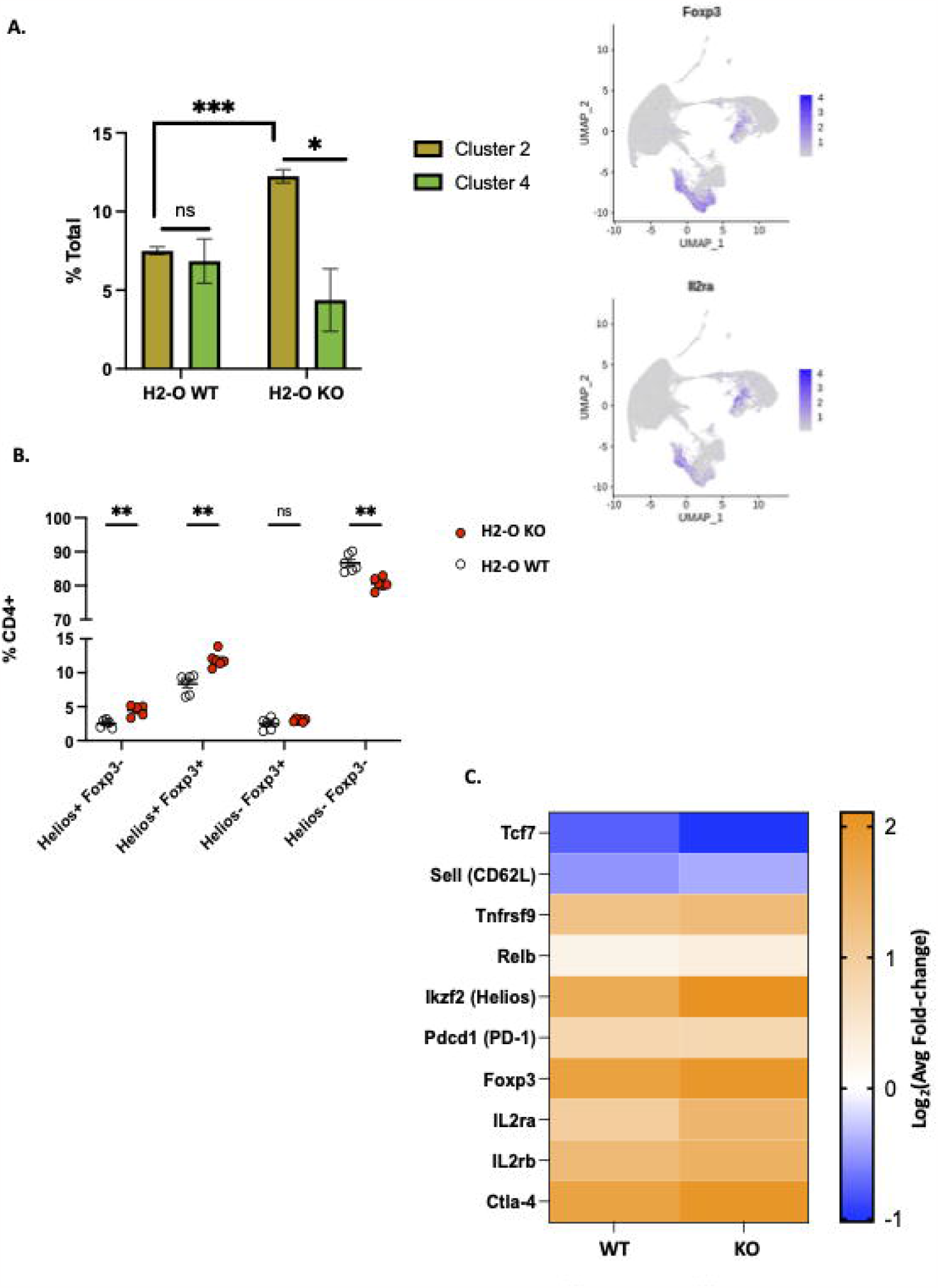
The H2-O KO Treg population has a more effector like phenotype. **A**.(Left) distribution of the two Treg populations identified by scRNA-seq clustering in either H2-O WT (left) or H2-O KO (right) samples. (Right) Expression of key Treg phenotypic markers: Foxp3 (Top) and CD25 [Il2ra] (Bottom) **B**.Subdivision of the splenic CD4+ T cell population by Foxp3 and Helios expression **C**.Comparison of cluster 2 upregulated genes (Log_2_Avg FC >0.5) in H2-O WT and H2-KO samples to a published (Miragaia *et al*.) splenic effector Treg genetic profile

In summary, both scRNA-seq and FACs analyses showed that H2-O KO mice have an increased frequency of both conventional and regulatory CD4 T cells in the spleens of unimmunized mice. Loss of thymic H2-O expression correlated with an increased tTreg population that was not from peripheral recirculation and had received increased TCR stimulation, which suggests an increased autoreactivity. Furthermore, increased activation of conventional splenic CD4 T cells is likely a driving factor for the increased effector-like Treg status that was identified by scRNA-seq analyses of H2-O-KO CD4 cells.

## Discussion

In this study, we have refined our understanding of how loss of H2-O alters CD4 T cell activity *in vivo*. Thymic analysis of medulla localized CD4 T cells pointed to a novel role for H2-O on CD4 T cells undergoing self-auditing. Importantly, lack of H2-O expression led to an increased frequency of both thymic and peripheral Tregs.

Tregs play a fundamental role in maintenance of homeostasis, hence a better understanding of the mechanism of their development is highly desirable and the subject of numerous studies (15, 19). Our studies here are the first attempt in demonstrating that H2-O plays a role in Treg development. While the exact mechanism by which thymic Treg cells are selected remains somewhat uncertain, recent studies support the idea that intermittent TCR signaling along with cytokine signaling drive Treg development (16). These findings and others (33, 34, 35, 36, 37) support a more avidity based model of Treg selection, in which alterations to the density of self-ligands present in the medulla will have differential effects on CD4 T cell selection. Intermittent TCR signaling as a driving force in Treg selection is very relevant to our understanding of how HLA-DO contributes to epitope selection during antigen processing. We have proposed that HLA-DO in complex with HLA-DM leads to better refinement of the epitopes from denatured antigens, promoting selection of the best fitting epitopes in the groove of MHC II molecules (11, 38). While this idea might appear in conflict with the multiple earlier interpretations of DO function (39, 40, 41), recent studies using human HLA-DO variants showed that certain variants enhanced DM activity (42).

Understanding of how DO works has been tumultuous over the past decades (5, 43, 44, 45, 46, 47, 48, 49). DO has been proposed to be an inhibitor of DM, or a helper of DM. Most reports acknowledge that understanding of DO requires further research. Our model that is based on studying kinetics of interactions of DO with DM and MHC class II using a variety of peptides of different binding characteristics is the most direct investigation of the biochemical mechanism of DO function. The most important evidence for our model come from data showing that DO or DO/DM binds to DR1 in a peptide receptive conformation: DO does not bind to the DR1 in complex with tightly bound peptides. Based on this model, when DO is present, higher affinity peptides are more likely to be presented. Conversely, in the absence of DO a larger percent of lower affinity epitopes are selected. Indeed, our in vivo peptide elution studies supports this biochemical mechanism as peptides eluted from H2-O KO mice expressing murine I-A^b^ (17) were of a lower general affinity. Accordingly, presentation of a larger portion of lower affinity peptides in the thymic medulla of H2-O KO mice is possible. Intermittent TCR signaling is typically generated by either pMHCII that are less stable (50, 51, 52, 53), or TCR/pMHC of lower affinity (54). Our findings here support the former. We associate increased Treg development as a consequence of the presentation of lower affinity peptides to self-auditing CD4 T cells.

Due to the continuous egress of mature CD4 T cells from the thymus, we also analyzed splenic CD4 T cells from unimmunized H2-O WT and H2-O KO mice. H2-O KO splenic CD4 T cells in unimmunized mice revealed an increased frequency of CD44hiCD69+ antigen experienced CD4 T cells, which suggests an increased level of antigen-specific TCR signaling in H2-O KO mice. As previously implied (55), a simple explanation for this activated state is incomplete thymic deletion in H2-O KO mice. While we were able to show that H2-O KO mice failed to delete specific CD4 clonal populations (17), detection of differences at the global levels did not show significant changes between the two genotypes. Nonetheless, we suggest that presentations of different arrays of self-antigens as well as their lower densities on thymic medulla leads to a less than optimal thymic deletion of self-reactive CD4 T cells and their routing to the periphery. Similar to activated CD4 T cells in the periphery, H2-O KO Treg cells also had a more effector-like phenotype, indicating enhanced Treg activation in unimmunized mice (32, 56).

In conclusion, our studies add a new dimension to our understanding of the role of H2-O in both CD4 T cell selection and activation. For the first time, we report on thymic negative selection in H2-O KO mice and demonstrate that loss of H2-O enhances thymic selection of regulatory T cells. Once in the periphery, an increased proportion of H2-O KO Tregs appear to be activated in a TCR dependent manner. These effector-like Tregs will likely help control increased basal CD4 T cell activation. However, the exact mechanism by which increased Treg cells are selected in the H2-O KO thymus remains to be determined. While we propose that alterations in medulla pMHCII-TCR avidity interactions could lead to enhanced Treg selection, increased presentation of self-antigens in a more tissue restricted antigenic (TRA) manner could also be possible (15).

## Methods

### Mice

Male and female C57BL/6J (H2-O WT, stock no: 000664), Female B6.PL-Thy1a/CyJ (stock no: 000406) were purchased from Jackson Laboratories and bred in house. Generation of the H2-O knock-out mice has been previously described (57) and mice bred in house. Unless otherwise stated analyzed mice were 6-8 weeks. All mouse procedures were approved by the Johns Hopkins University Animal Care and Use Committee and were following relevant ethical regulations.

### Antibodies/reagents

#### Flow Cytometry

anti-CD3e (17A2), anti-CD4 (RM4-5), anti-CD5 (53-7.3), anti-CD8α (53-6.7), CD11c (N418), anti-CD19 (ID3), anti-CD25 (PC61), anti-CD44 (IM7), anti-CD45R/B220 (RA3-6B2), anti-CD62L (MEL-14), anti-CD69 (H1.2F3), anti-CD197/CCR7 (4B12), anti-NK1.1 (PK136), anti-TCRβ (H57-597), anti-TCRγδ (GL3), anti-Cleaved Caspase 3 (D3E9) Cell signaling (Danvers, MA); Foxp3 (150D), anti-Helios (22F6), anti-Nur77 (12.14), Fixable viability dye eFluor 780 (eBioscience).

#### FACs Sorting

anti-CD3e, anti-CD4, anti-CD19, anti-NK1.1, Propidium iodide. Briefly, 30,000 live CD4 T cells [Gating: CD3+, CD4, CD19-, NK1.1-, PI-] were FACs sorted from the spleens of 3 unimmunized 10 week-old H2-O WT and 3 unimmunized 10 week-old unimmunized H2-O KO female mice.

### Cell staining

For cleaved caspase 3 staining (18) homogenized mice thymocyte cells were stained with anti-CCR7/CD197 at a final dilution of 1:50 for 30 min at 37°C prior to additional surface stains. Following surface staining, cells were fixed with Cytofix/Cytoperm (BD Biosciences) for 20 min at 4°C. Cells were then washed with Perm/Wash buffer (BD Biosciences) twice. Cells were stained with anti–cleaved caspase 3 at a 1:50 dilution at 23°C for 30 min.

For transcription factor staining, cells were incubated with surface antibody at 4°C for 20 min, permeabilized at 4°C for 30 min using a Foxp3/Transcription factor buffer set (Invitrogen, ThermoFisher Scientific), and then stained with anti–Foxp3 and/or anti-Helios at 23°C for 30 min.

### Adoptive Cell Transfer

Naïve CD4 T cells were isolated from the pooled spleens and LNs of 4 week old Thy1.1 expressing C57BL/6J mice using the EasySep Mouse Naive CD4 T cell Isolation Kit (StemCell Technologies). Isolated cells were stained with eFluor 450 viability dye (eBiosciences) according to manufactures directions. 3x10^6^ dye labeled Thy1.1 cells were IV injected into 10 week old H2-O WT or H2-O KO hosts. Transferred cells were recovered 7 days post-transfer.

### scRNA-Sequencing

#### Library & sequencing

The samples were prepared using the 10x Genomics Chromium Next GEM Single Cell 5’ Library and Gel Bead Kit v1.1, Chromium Next GEM Chip and Dual Index Kit TN Set A. They were run on the Illumina NovaSeq6000 with a run configuration of 28bp x 10bp x 10bp x 91bp.

#### Analysis

Analyses of T-cell scRNA-seq were performed with the package Seurat (58), as follows. Data was filtered to remove cells with low gene count (<200), large number of UMIs (>12,000) and high (>5%) fraction of mitochondrial reads. Expression levels of genes were log-normalized, and the most variable 2000 genes were selected for linear dimensionality reduction with Principal Component Analysis (PCA). The first 15 principal components were then used to performed unsupervised clustering using the Seurat SNN clustering package, with a resolution of 0.2. To identify cell types, potential markers for each cluster were calculated as the set of genes significantly differentially expressed in each cluster compared to all others, using the function FindAllMarkers, and by searching existing literature and marker databases. Lastly, differentially expressed genes for each cell type between conditions were determined using the function FindMarkers with the default function (bimod).

Defining clusters – Genes with an average fold-change (avgFC) >0.5 and an adjusted p-value<0.05 were run against CD4 T cell data sets available on the Immunological Genome Project (https://www.immgen.org/) using the “My Geneset” data browser function.

### Statistical testing

GraphPad Prism was used for all statistical analyses. A standard Student T-test was used for estimation of statistical significance. Data is shown as mean ± SEM. *p<0.05, **p<0.01,***p<0.001, ****p<0.0001.

## Supporting information

Individual Replicate Information

Cells per cluster

## Acknowledgments

Authors thank Drs Lars Karlsson and Peter Jensen for the gift of H2-O KO mice. We also thank Dr. Liliana D. Florea, and Corina Antonescu from the Computational Biology Computing Core at Johns Hopkins School of Medicine for help with scRNA-Seq analysis. FACs analysis was performed in the BD Immunology and Flow Cytometry Laboratory at the Johns Hopkins School of Public Health. Supported by grants from NIAID, R01AI063764, R21AI101987, and R01AI120634, to SS-N.

## Figure Legends

**Supplemental Figure 1: Gating strategy used for detection of Auditing vs Clonally Deleted Medulla CD4 T cells** Red arrows indicate how Auditing vs Clonal Deletion was identified in 6-week-old H2-O WT and H2-O KO thymi.

Blue arrows indicate additional gating corresponding to Figure 1a.

Black Arrows indicate additional gating corresponding to Figure 1b.

**Supplemental Figure 2: The frequency of auditing Mature 1 CD4 T cells was increased in H2-O KO thymi A**. Percentage of auditing (Caspase-3 negative) CD4 T cells in the Semi-mature (CD69+ MHC-I neg), Mature 1 (CD69+ MHC-1+), and Mature 2 (CD69-MHC-I+) in H2-O WT (white) and H2-O KO (red) mice N= 18 mice per group

**B**. Percentage of clonally deleted (Caspase-3+) CD4 T cells in the Semi-mature (CD69+ MHC-I neg), Mature 1 (CD69+ MHC-1+), and Mature 2 (CD69-MHC-I+) in H2-O WT (white) and H2-O KO (red) mice. N= 18 mice per group Statistical Testing: unpaired student T-test

**Supplemental Figure 3: Positive Selection is not affected by loss of H2-O**

A. Representative plots showing CD4, CD8 and Double Positive (DP) percentages in H2-O WT (Top) and H2-O KO (Bottom)

B. Summary plots of the single-positive CD4 (Top) and CD8 (Bottom) percentages from 5 repeat experiments.

C. (Left) representative flow plots showing the “Signaled” vs “Non-signaled” thymocytes in H2-O WT (Top) or H2-O KO (Bottom). (Right) Summary plots of >5 repeat experiments.

D. Frequency of signaled CCR7+ medulla CD4 T cells in H2-O WT (white) and H2-O KO (red) mice

**Supplemental Figure 4:** Top: breakdown of H2-O KO (right) and H2-O WT (Left) Seurat clustering

Bottom: Seurat clustering for each individual biological replicate

## Notes

### Competing Interest Statement

The authors have declared no competing interest.

### Summary of Updates

Figures 1 and 2 have been updated for clarity and supplemental Figures have been updated to include gating strategies

