## Supplementary figures and images for "H2-O deficiency promotes regulatory T cell differentiation and CD4 T cell hyperactivity"

### Supplemental Figure 1.tiff

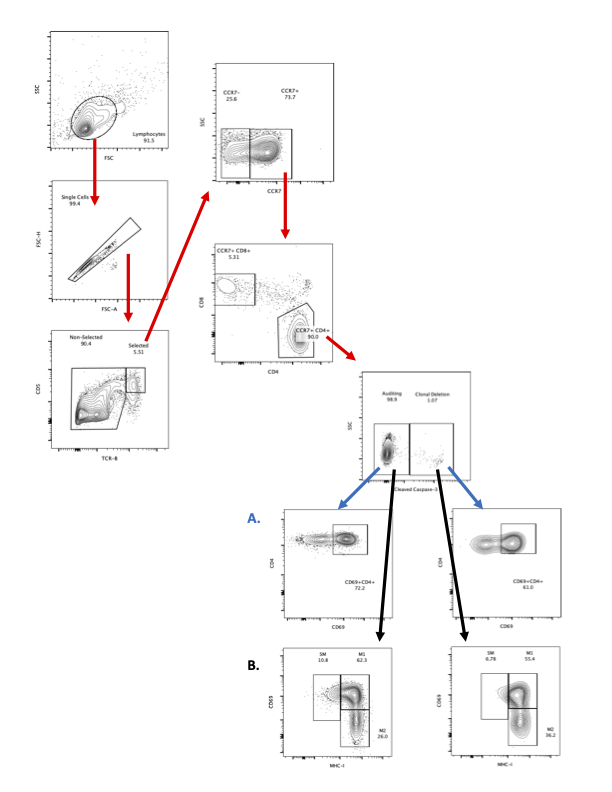

### Supplemental Figure 2.tiff

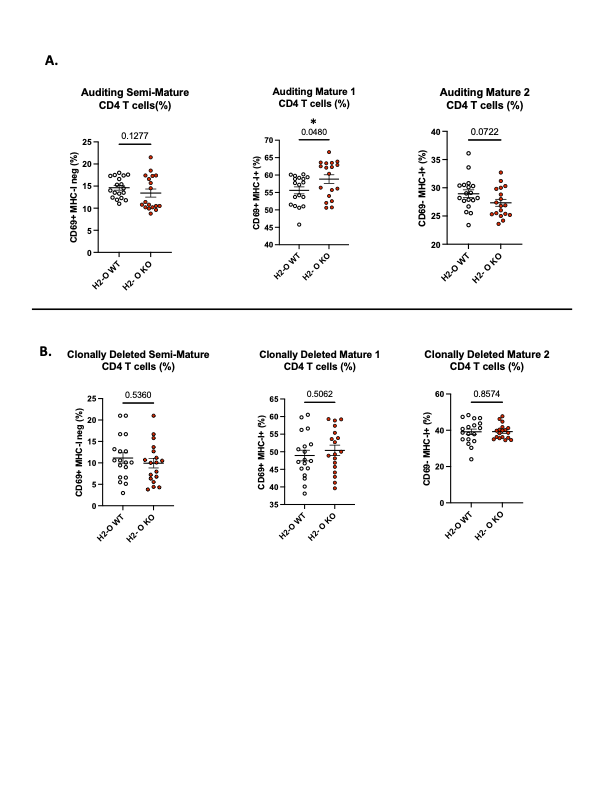

### Supplemental Figure 3.tiff

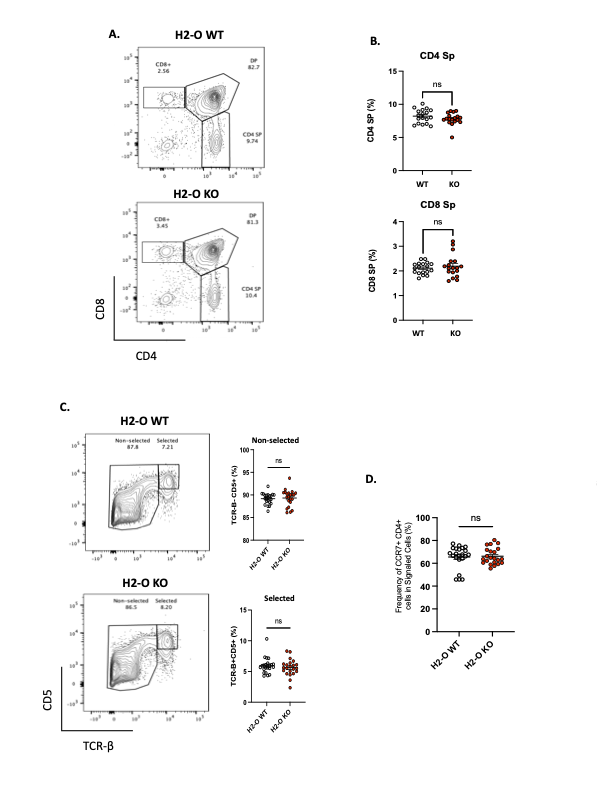

### Supplemental Figure 4.tiff

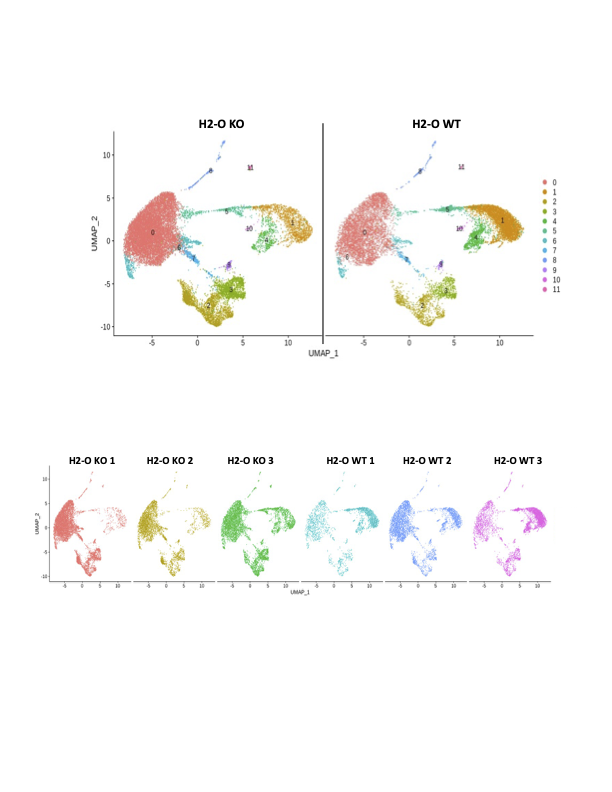
